# Metagenome-Assembled Genome Binning Methods with Short Reads Disproportionately Fail for Plasmids and Genomic Islands

**DOI:** 10.1101/2020.03.31.997171

**Authors:** Finlay Maguire, Baofeng Jia, Kristen Gray, Wing Yin Venus Lau, Robert G. Beiko, Fiona S.L. Brinkman

**Affiliations:** Faculty of Computer Science, Dalhousie University; Department of Molecular Biology and Biochemistry, Simon Fraser University

## Abstract

Metagenomic methods are an important tool in the life sciences, as they enable simultaneous characterisation of all microbes in a community without time-consuming and bias-inducing culturing. Metagenome-assembled genome (MAG) binning methods have emerged as a promising approach to recover individual genomes from metagenomic data. However, MAG binning has not been well assessed for its ability to recover mobile genetic elements (MGEs), such as plasmids and genomic islands (GIs), that have very high clinical/agricultural/environmental importance. Certain antimicrobial resistance (AMR) genes and virulence factor (VF) genes are noted to be disproportionately associated with MGEs, making studying their transmission a public health priority. However, the variable copy number and sequence composition of MGEs relative to the majority of the host genome makes them potentially problematic for MAG binning methods. To systematically investigate this, we simulated a low-complexity metagenome comprising 30 GI-rich and plasmid-containing bacterial genomes. MAGs were then recovered using 12 current prediction pipelines and evaluated for recovery of MGE-associated AMR/VF genes. Here we show that while 82-94% of chromosomes could be correctly recovered and binned, only 38-44% of GIs were recovered and, even more notably, only 1-29% of plasmid sequences were found. Most strikingly, no plasmid-borne VF or AMR genes were recovered and within GIs, only between 0-45% of AMR or VF genes were identified. We conclude that short-read MAGs are largely ineffective for the analysis of mobile genes, including those of public-health importance like AMR and VF genes. We propose that microbiome researchers should instead primarily utilise unassembled short reads and/or long-read approaches to more accurately analyse metagenomic data

## Main

Metagenomics, the sequencing of DNA from within an environmental sample, is widely used to characterise the functional potential and identity of microbial communities [1,2]. These approaches have been instrumental in developing our understanding of the distribution and evolutionary history of AMR genes [3,4,5], as well as tracking pathogen outbreaks [6]. Although long-read DNA technologies (e.g., Oxford Nanopore [7], PacBio [8]) are now being used for metagenomic sequencing [9,10], high-throughput sequencing of relatively short reads (150-250bp) in platforms such as the Illumina MiSeq still dominates metagenomics. These reads can be directly analysed using reference databases and a variety of homology search tools (e.g., [11,12,13,14]). Since these reads are shorter than most genes, however, read-based methods provide very little information about their genomic organisation. This lack of contextual information is particularly problematic in the study of AMR genes and VFs as the genomic context plays a role in function [15], selective pressures [16], and likelihood of lateral gene transfer (LGT) [17].

Sequence assembly using specialised metagenomic de Bruijn graph assemblers (e.g., metaSPAdes [18], IDBA-UD [19], and megahit [20]) is often used to try to recover information about genomic context [21]. To disentangle the resulting mix of assembled fragments, there has been a move to group these contigs based on the idea that those from the same source genome will have similar relative abundance and sequence composition [22]. These resulting groups or “bins” are known as metagenome-assembled genomes (MAGs). A range of tools have been released to perform this binning including CONCOCT [23], MetaBAT 2 [24], MaxBin 2 [25], and a tool which combines their predictions: DAS Tool [26]. These MAG binning methods have been used in unveiling previously uncharacterised genomic diversity [27,28,29], but metagenomic assembly and binning results in the loss of some information. This compounded data loss means as little as 24.2-36.4% of reads [30,31] and ~23% of genomes [31] are successfully assembled and binned in some metagenomic analyses. The Critical Assessment of Metagenome Interpretation (CAMI) challenge’s (https://data.cami-challenge.org/) Assessment of Metagenome BinnERs (AMBER) [32] benchmarks different MAG recovery methods in terms of global completeness and bin purity. However, to the best of our knowledge, there has not been a specific assessment of MAG-based recovery of mobile genetic elements (MGEs) like genomic islands (GIs) and plasmids, despite their health and research importance.

Genomic islands (GIs) are clusters of genes that are known or predicted to have been acquired through LGT events. GIs can arise following the integration of MGEs, such as integrons, transposons, integrative and conjugative elements (ICEs) and prophages (integrated phages) [33,34]. They are of high interest since VFs are disproportionately associated with mobile sequences [35] as well as certain AMR genes [36,37]. GIs often have differing nucleotide composition compared to the rest of the genome [33], a trait exploited by GI prediction tools such as SIGI-HMM [38], IslandPath-DIMOB [39], and integrative tools like IslandViewer [40]. GIs may also exist as multiple copies within a genome [41] leading to potential assembly difficulties and biases in the calculation of coverage statistics.

Plasmids are circular or linear extrachromosomal self-replicating pieces of DNA with variable copy numbers and repetitive sequences [42,43]. Similar to GIs, the sequence composition of plasmids are often markedly different from the genome with which they are associated [44,45]. Plasmids are also of high interest as a major source of the lateral dissemination of AMR genes throughout microbial ecosystems [36,46].

These varying composition and relative abundance features mean that GIs and plasmids pose significant challenges in MAG recovery. As these MGEs are key to the function and spread of pathogenic traits such as AMR and virulence, and with MAG approaches becoming increasingly popular within microbial and public-health research, it is both timely and vital that we assess the impact of metagenome assembly and binning on the recovery of these elements. Therefore, to address this issue we performed an analysis of GI and plasmid (and associated AMR/VF genes) recovery accuracy across a set of 12 state-of-the-art methods for short-read metagenome assemblies. We show that short-read MAG-based analyses are not suitable for the study of mobile sequences.

## Results

### Recovery of Genomic Elements

#### Chromosomes

The overall ability of MAG methods to recover the original chromosomal source genomes varied widely. We considered the “identity” of a given MAG bin to be that of the genome that comprises the largest proportion of sequence within that bin. In other words if a bin is identifìably 70% species A and 30% species B we consider that to be a bin of species A. Ideally, we wish to generate a single bin for each source genome consisting of the entire genome and no contigs from other genomes. Some genomes are cleanly and accurately binned regardless of the assembler and binning method used (see Fig. 1). Specifically, greater than 90% of *Streptomyces parvulus* (minimum 91.8%) and *Clostridium baratii* (minimum 96.4%) chromosomes are represented in individual bins across all methods. However, no other genomes were consistently recovered at >30% chromosomal coverage across methods. The three *Streptococcus* genomes were particularly problematic with the best recovery for each ranging from 1.7% to 47.49%. Contrary to what might be expected, the number of close relatives to a given genome in the metagenome did not clearly affect the MAG coverage (Fig. S1).

**Figure 1:**
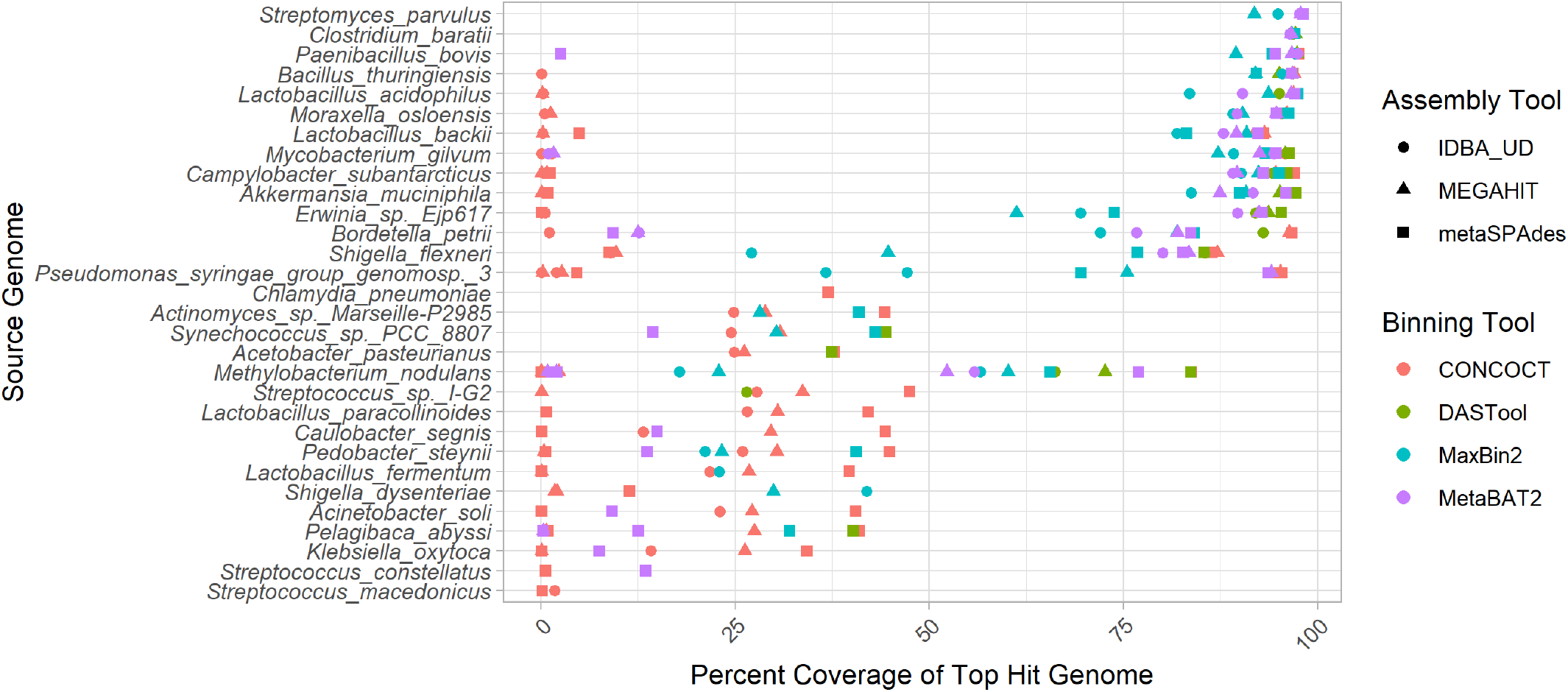
Top genome coverage for input genomes across MAG binners. Each dot represents the coverage of a specified genome when it comprised the plurality of the sequences in a bin. If a genome did not form the plurality of any bin for a specific binner-assembler pair no dot was plotted for that genome and binner-assembler. The binning tool is indicated by the colour of the dot as per the legend. Genomes such as *Clostridium baratti* were accurately recovered across all binner-assembler combinations whereas genomes such as *Streptococcus macedonicus* were systematically poorly recovered.

In terms of the impact of different metagenome assemblers, megahit resulted in the highest median chromosomal coverage across all binners (81.9%) with metaSPAdes performing worst (76.8%) (Fig. 2 A). In terms of binning tools, CONCOCT performed very poorly with a median 26% coverage for top hit per bin, followed by maxbin2 (83.1%), and MetaBAT2 (88.5%). It is perhaps unsurprising that the best-performing binner in terms of bin top hit coverage was the metabinner DASTool that combines predictions from the other 3 binners (94.3% median top hit chromosome coverage per bin; (Fig. 2 A)).

**Figure 2:**
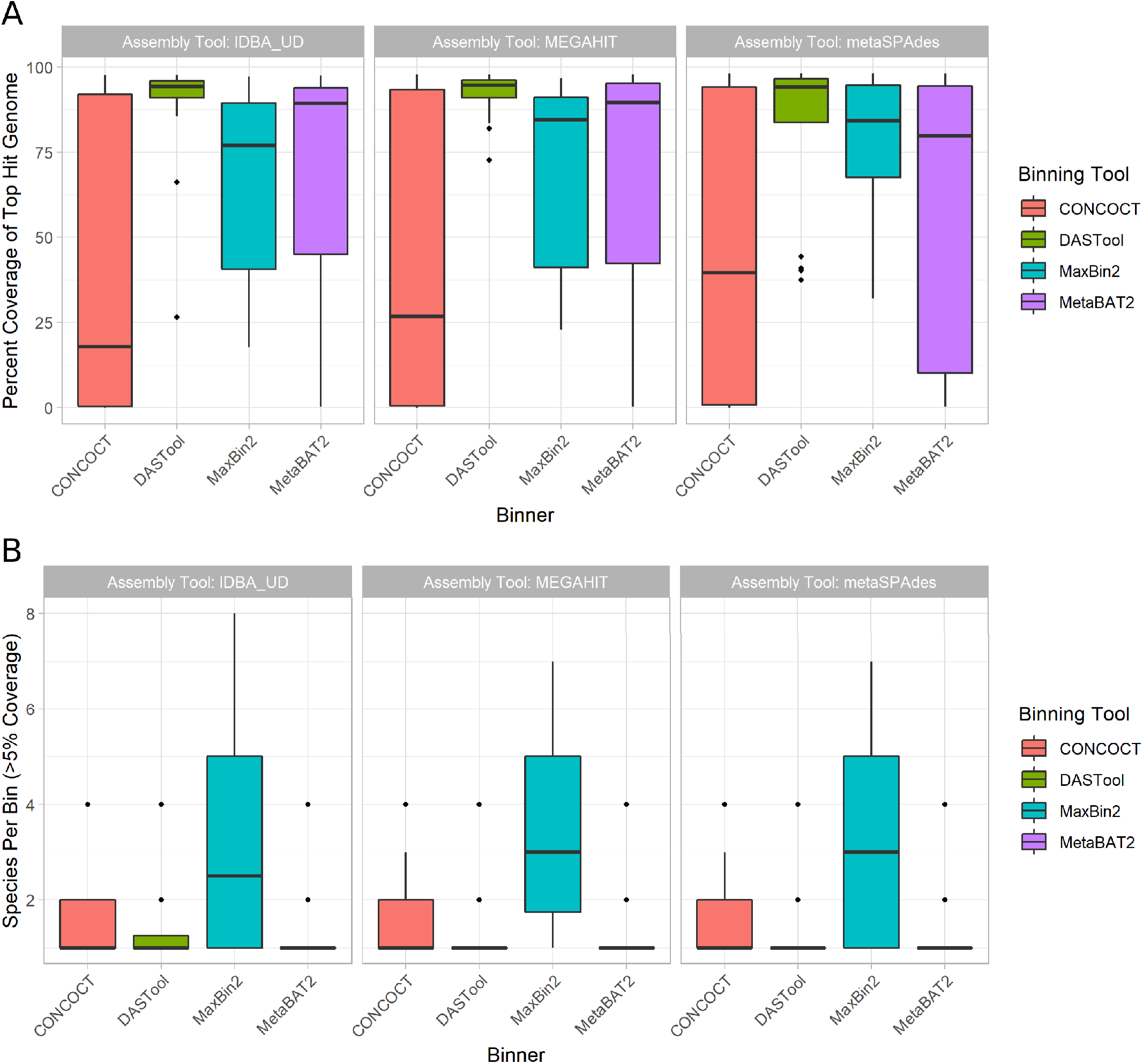
Overall binning performance for every combination of metagenome assembler (as indicated by pane titles) and MAG binning tool (x-axis and legend colours). Diamonds in the plots represent outliers (greater or lower than the interquartile range marked by the error bars) and the boxes represent the lower quartile, median, and upper quartile respectively. **(A)** Chromosomal coverage of the most prevalent genome in each bin across binners and metagenome assemblies. Of the 3 assemblers, megahit resulted in the highest median chromosomal coverage (y-axis) across all binners (colored bars) at 81.9% with metaSPAdes performing the worst (76.8%). Of the 4 binners, CONCOCT (red) performed poorly with a median coverage, followed by maxbin2 (blue), MetaBAT2 (purple) and DASTool (green) performing the best. **(B)** Distribution of bin purity across assemblers and binners. The total number of genomes present in a bin at >5% coverage (y-axis) was largely equivalent across assemblers (x-axis). For the binning tools, maxbin2 (blue) produced nearly twice as many bins containing multiple species compared to CONCOCT (red), MetaBAT2 (purple) and DASTool (green), which all produced chimeric bins at roughly the same rate.

Bin purity, i.e. the number of genomes present in a bin at >5% coverage, was largely equivalent across assemblers, with a very marginally higher purity for IDBA. Across binning tools maxbin2 proved an exception with nearly twice as many bins containing multiple species as the next binner (Fig. 2 B). The remaining binning tools were largely equivalent, producing chimeric bins at approximately the same rates. Unlike coverage, purity was strongly affected by the number of close relatives in the metagenome to a given input genome. Specifically, the closer the nearest relative the less pure the bin (Fig. S2).

#### Plasmids

Regardless of method, a very small proportion of plasmids were correctly grouped in the bin that was principally composed of chromosomal contigs from the same source genome. Specifically, between 1.5% (IDBA-UD assembly with DASTool bins) and 29.2% (metaSPAdes with CONCOCT bins) were correctly binned at over 50% coverage. In terms of metagenome assembly, metaSPAdes was by far the most successful assembler at assembling plasmids with 66.2% of plasmids identifiable at greater than 50% coverage. IDBA-UD performed worst with 17.1% of plasmids recovered, and megahit recovered 36.9%. If the plasmid was successfully assembled, it was, with one exception, placed in a MAG bin by maxbin2 and CONCOCT, although a much smaller fraction were correctly binned (typically less than 1/3rd). Interestingly, the MetaBAT2 and DASTool binners were more conservative in assigning plasmid contigs to bins; of those assigned to bins, nearly all were correctly binned (Fig. 3).

**Figure 3:**
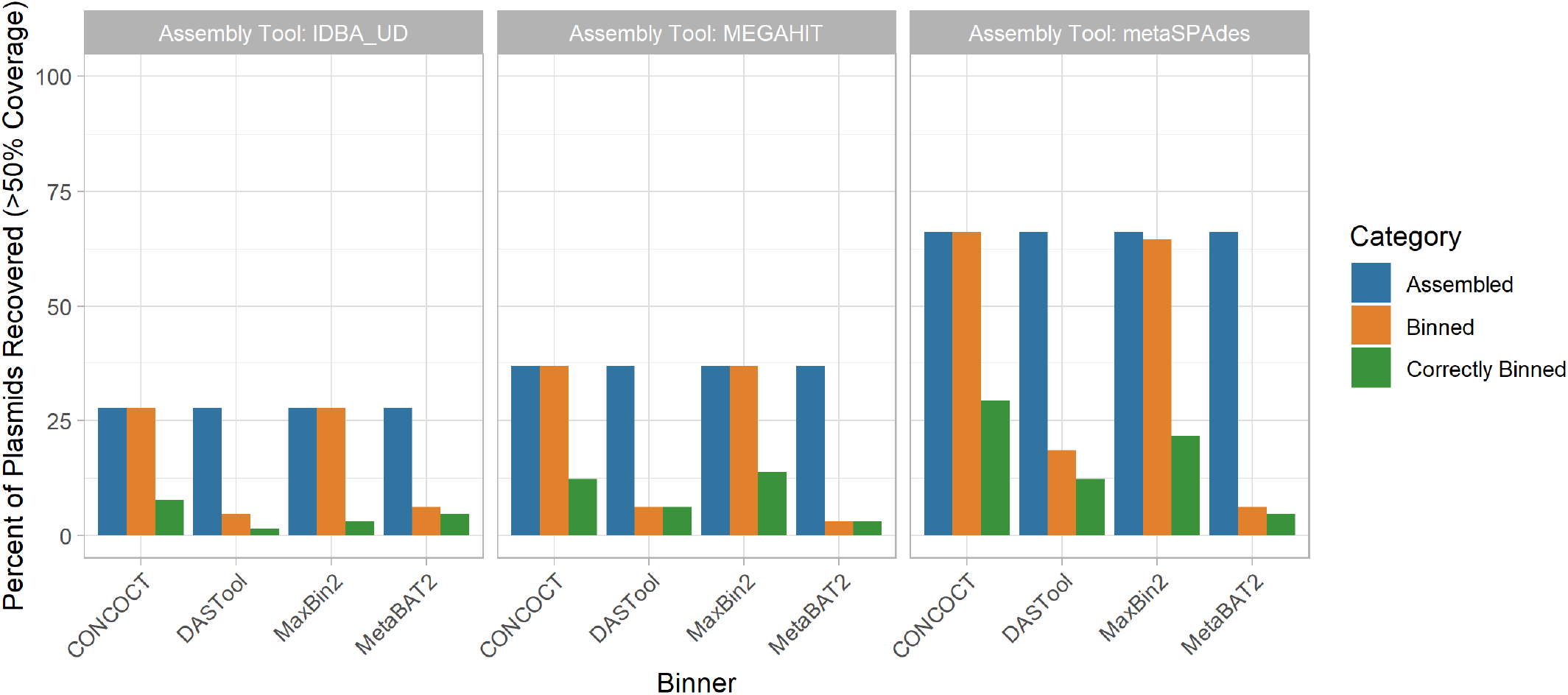
The performance of metagenomic assembly and binning to recover plasmid sequences. Each plot represents a different metagenome assembler, with the groups of bars along the x-axes showing the plasmid recovery performance of each binning tool when applied to the assemblies produced by that tool. For each of these 12 assembler-binner-pair-produced MAGs the grouped bars from left to right show the percentage of plasmids assembled, assigned to any bin, and binned with the correct chromosomes. These stages of the evaluation are indicated by the bar colours as per the legend. Across all tools the assembly process resulted in the largest loss of plasmid sequences and only a small proportion of the assembled plasmids were correctly binned.

#### Genomic Islands

GIs displayed a similar pattern of assembly and correct binning performance as plasmids (Fig. 4). Assembly of GIs with >50% coverage was consistently poor (37.8-44.1%) with metaSPAdes outperforming the other two assembly approaches. For the CONCOCT and maxbin2 binning tools, all GIs that were assembled were assigned to a bin, although the proportion of binned GIs that were correctly binned was lower than for DASTool and MetaBAT2. DASTool, MetaBAT2 and CONCOCT did not display the same precipitous drop between those assembled and those correctly binned as was observed for plasmids. In terms of overall correct binning with the chromosomes from the same genome the metaSPAdes assembly with CONCOCT (44.1%) and maxbin2 (43.3%) binners performed best.

**Figure 4:**
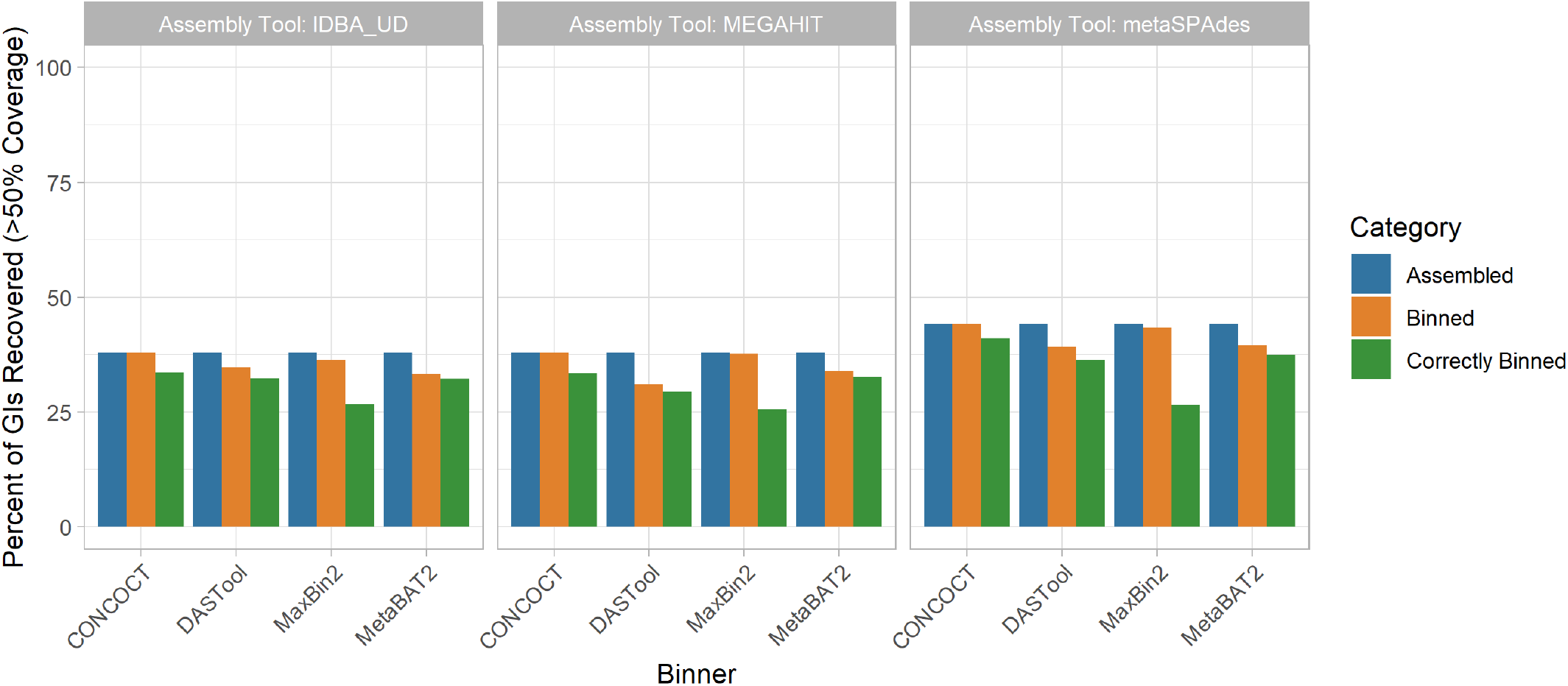
Impact of metagenomic assembly and MAG binning on recovery of GIs. GIs were recovered in a similarly poor fashion to plasmids. Generally, <40% were correctly assigned to the same bin majorly comprised of chromosomal contigs from the same source genome regardless of binning (x-axis) and assembly (panel) methods at >50% coverage. metaSPAdes performed the best at assembling GIs (blue). Maxbin2 and CONCOCT placed GIs in a bin majority of the time (orange) however a very small fraction was correctly binned (green). Generally, GIs were correctly binned better than plasmids with DASTool, MetaBAT2 and CONCOCT.

#### AMR Genes

The recovery of AMR genes in MAGs was poor with only ~49-55% of all AMR genes predicted in our reference genomes regardless of the assembly tool used, and metaSPAdes performing marginally better than other assemblers (Fig. 5 A). Binning the contigs resulted in a ~1-15% loss in AMR gene recovery with the CONCOCT-metaSPAdes pair performing best at only 1 % loss and DASTool-megahit performing the worst at 15% reduction of AMR genes recovered. Overall, only 24% - 40% of all AMR genes were correctly binned. This was lowest with the maxbin2-IDBA-UDA pair (24%) and highest in the CONCOCT-metaSPAdes pipe (40%).

**Figure 5:**
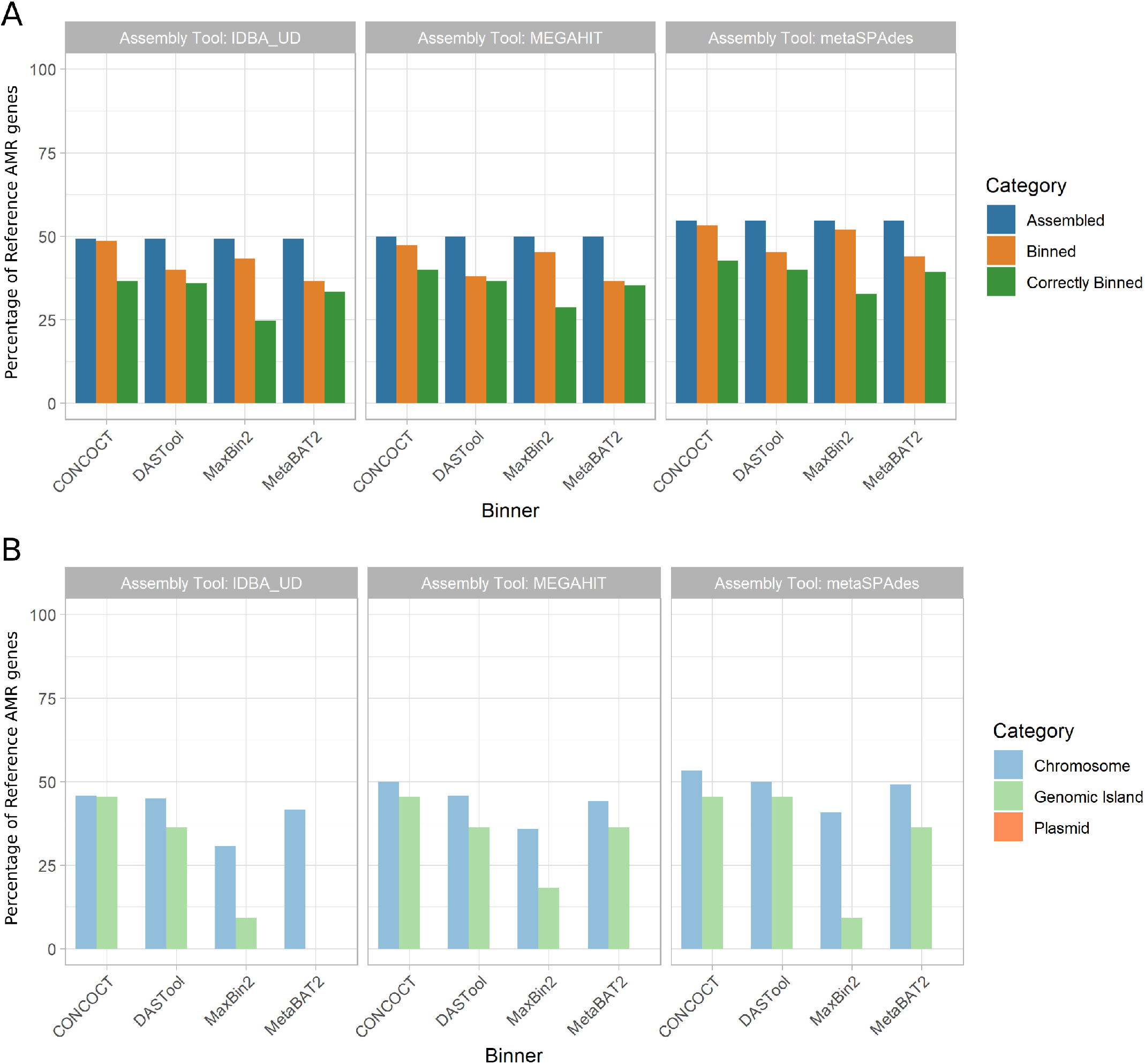
Recovery of AMR genes across assemblers, binners, and genomic context. **(A)** The proportion of reference AMR genes recovered (y-axis) was largely similar across assembly tools (panels as indicated by title) at roughly 50% with metaSPAdes performing marginally better overall. Binning tools (x-axis) resulted in a small reduction in AMR genes recovered (orange), however only 24-40% of all AMR genes were correctly binned (green). metaSPAdes-CONCOCT was the best performing MAG binning pipeline. **(B)** Percent of correctly binned AMR genes recovered by genomic context. MAG methods were best at recovering chromosomally located AMR genes (light blue) regardless of metagenomic assembler or binning tool used. Recovery of AMR genes in GIs showed a bigger variation between tools (light green). None of the 12 evaluated MAG recovery methods were able to recover plasmid located AMR genes.

Moreover, focusing on only the AMR genes that were correctly binned (Fig. 5 B) we can evaluate the impact of different genomic contexts (i.e. chromosomal, plasmid, GI). Across all methods only 30%-53% of all chromosomally located AMR genes (n=120), 0-45% of GI located AMR genes (n=11) and none of the plasmid-localised AMR genes (n=20) were correctly binned.

#### Virulence Factor Genes

We also examined the impact of MAG approaches on recovery of virulence factor (VF) genes as identified using the Virulence Factor Database (VFDB). We saw a similar trend as AMR genes (Fig. 6 A). Between 56% and 64% of VFs were identifiable in the metagenomic assemblies (with megahit recovering the greatest proportion). The binning process further reduced the number of recovered VFs by 4-26% with DASTool-megahit performing the worst (26% reduction) and CONCOCT-metaSPAdes performing the best (4% reduction). Unlike AMR genes, the majority of VF genes assigned to a bin were assigned to the correct bin (i.e. that bin largely made up of contigs from the same input genome). Overall, CONCOCT-metaSPAdes again performed best with 43% of all VFs correctly assigned.

**Figure 6:**
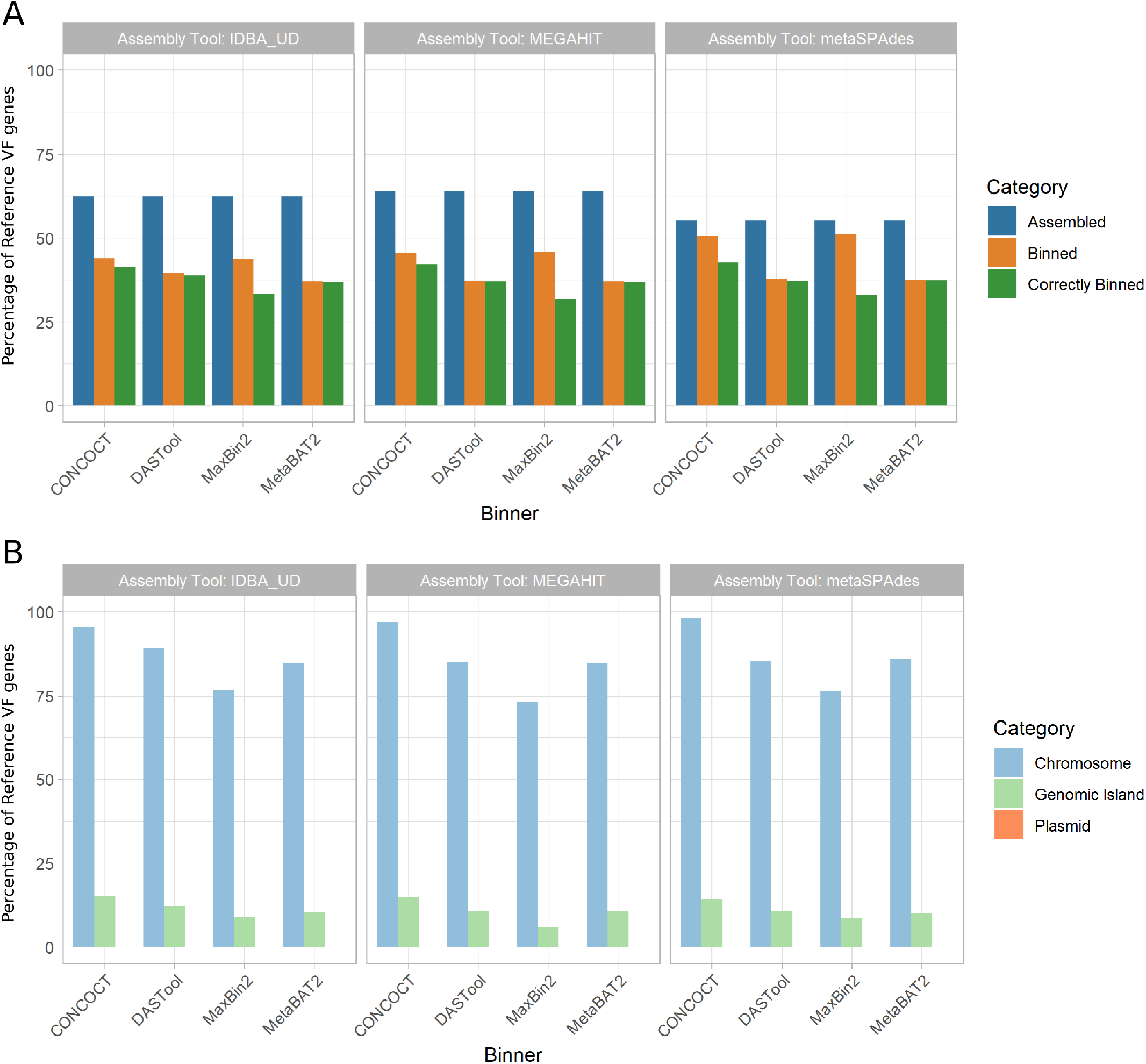
Recovery of VF genes across assemblers, binners, and genomic context. **(A)** Percent of reference virulence factor (VF) genes recovered across assemblers and binners. The proportion of reference VF genes recovered (y-axis) exhibited a similar trend as AMR genes. Recovery was greatest after the assembling stage (blue), with megahit performing best. Binning tools resulted in a larger reduction in VF genes recovered (orange) compared to AMR genes. However, in the majority of cases, VF genes that are binned are correctly binned (green). metaSPAdes-CONCOCT was again the best performing pair. **(B)** Percent of correctly binned VF genes recovered in each genomic region. Metagenome assembled genomes (MAGs) were again best at recovering chromosomally located VF genes (light blue), able to correctly bin majority of chromosomally located VFs. GIs recovered again performed very poorly (light green) and again none of the plasmid located AMR genes (orange) was correctly binned.

As with AMR genes, the genomic context (chromosome, plasmid, GI) of a given VF largely determined how well it was binned (Fig. 6 B). The majority (73%-98%) of all chromosomally located VF genes (n=757) were correctly binned. However, 0-16% of GI-localised VF genes (n=809) and again none of the plasmid-associated VF genes (n=3) were recovered across all 12 MAG pipelines.

### Comparisons of Rates of Loss

We combined the performance metrics for Figs. 3, 4, 5, and 6 to compare the rates of loss of different components (see Fig. S4). This highlighted that genomic components (GIs and plasmids) and plasmids in particular are lost at a disproportionately higher rate than individual gene types during MAG recovery.

## Discussion

In this paper, we evaluated the ability of metagenome-assembled genome (MAG) binning methods to correctly recover mobile genetic elements (MGEs; i.e. GIs and plasmids) from metagenomic samples. Overall, chromosomal sequences were binned well (up to 94.3% coverage, with perfect bin purity using megahit-DASTool) however closely related genomes were consistently cross-contaminated with other sequences (e.g. *Streptococcus* species in Fig. S1, S2). Given the importance of MGEs in the function and spread of pathogenic traits, it is particularly noteworthy that regardless of MAG binning method, plasmids and GIs were disproportionately lost compared to core chromosomal regions. At best (with metaSPAdes and CONCOCT) 29.2% of plasmids and 44.1% of GIs were identifiable at >50% coverage in the correct bin (i.e. grouped with a bin that was mostly made up of contigs from the same genome). While some MGEs were likely recovered in more partial forms (<50% coverage), use of these by researchers interested in selective pressures and lateral gene transfer could lead to inaccurate inferences. This poor result is congruent with the intuition that the divergent compositional features and repetitive nature of these MGEs is problematic for MAG methods. The particularly poor plasmid binning performance is likely attributable to the known difficulties in assembly of plasmids from short-read data [47]. Therefore, binning efficiency might improve with use of long-read sequencing or assembly methods optimised for recovering plasmids [47] (such as SCAPP [48]). Despite its lower effective sequencing depth and higher error rates, incorporating long-read sequencing has been shown to improve overall MAG binning [49] and facilitate metagenomic characterisation of plasmids [50]. Further research is needed to fully characterise the performance of different long-read protocols on the accuracy of recovering MGEs in metagenomic samples.

With the growing use of MAG methods in infectious disease research (e.g., [51,52,53,54,55]) and the public-health importance of the LGT of AMR and VF genes, we also specifically evaluated the binning of these gene classes. The majority of these genes were correctly assembled across assemblers but were either not assigned or incorrectly assigned to MAG bins during binning. At best across all binners, 40% of all AMR genes and ~63% of VF genes (CONCOCT-metaSPAdes) present in the reference genomes were assigned to the correct MAG. While a majority of chromosomally located VF genes (73-98%) and AMR genes (53%) were binned correctly, only 16% of GI VFs (n=809), 45% of GI AMR genes (n=11), and not a single plasmid associated VF (n=3) or AMR gene (n=20) were correctly binned. This included critical high-threat MGE-associated AMR genes such as the KPC and OXA carbapenemases. One potential caveat of this is that some AMR genes and VFs may no longer be detectable in MAGs due to issues with ORF prediction (see suppl. discussion & Fig. S3). Previous studies have observed that ORF predictions in draft genomes are more fragmented, which can lead to downstream over- or under-annotation with functional labels depending on the approach used [56]. Although not yet developed, methods that combine the assembly/binning pipelines tested here with read-based inference would give a better sense of which functions are potentially being missed by the MAG reconstructions.

Our simulated metagenomic community comprised 30 distinct bacterial genomes with varying degrees of relatedness. While this diversity can be representative of certain clinical samples [57,58,59], other environments with relevance to public health such as the human gut, soil, and livestock can have 100-1000s of species [60,61,62,63]. Consequently our analysis likely overrepresents the effectiveness of the methods tested in a public-health setting. Metagenomic simulation is also unlikely to perfectly represent the noise and biases in real metagenomic sequencing but it does provide the ground-truth necessary for evaluation [32,64]. This simulation approach, combined with the development of an MGE/AMR-focused mock metagenome (similarly to the mockrobiota initiative [65]), could provide a key resource to develop and validate new binning approaches and different sequencing strategies.

## Conclusions

This study has shown that MAG-based approaches provide a useful tool to study a bacterial species’ core chromosomal elements, but have severe limitations in the recovery of MGEs. The majority of these MGEs will either fail to be assembled or be incorrectly binned. The consequence of this is the disproportionate loss of key public-health MGE-associated VFs and AMR genes. As many of these clinically relevant genes have a high propensity for lateral gene transfer between unrelated bacteria [35,36] it is critical to highlight that MAG approaches alone are insufficient to thoroughly profile them. Within public-health metagenomic research it is vital we utilise MAGs in conjunction with other methods (e.g. targeted AMR [66], long-read sequencing, plasmid specialised assembly approaches [48], and read-based sequence homology search [11]) before drawing biological or epidemiological conclusions.

## Methods

In keeping with FAIR principles (Findable, Accessible, Interoperable, Reusable data), all analyses presented in this paper can be reproduced and inspected with the associated github repository github.com/fmaguire/MAG_gi_plasmid_analysis and data repository osf.io/nrejs/.

### Metagenome Simulation

Thirty RefSeq genomes were selected using IslandPath-DIMOB [39] GI prediction data collated into the IslandViewer database www.pathogenomics.sfu.ca/islandviewer [40] (Supplemental Table 1). The selected genomes and associated plasmids (listed in Supplemental Table 2 and deposited at osf.io/nrejs/ under “data/sequences”) were manually selected to satisfy the following criteria:

1. 10 genomes with 1-10 plasmids.
2. 10 genomes with >10% of chromosomal DNA predicted to reside in GIs.
3. 10 genomes with <1% of chromosomal DNA predicted to reside in GIs.

In accordance with the recommendation in the CAMI challenge [67] the genomes were randomly assigned a relative abundance following a log-normal distribution (μ = 1, σ = 2). Plasmid copy number estimates could not be accurately found for all organisms. Therefore, plasmids were randomly assigned a copy number regime: low (1-20), medium (20-100), or high (500-1000) at a 2:1:1 rate. Within each regime, the exact copy number was selected using an appropriately scaled gamma distribution (α = 4, β = 1) truncated to the regime range.

Finally, the effective plasmid relative abundance was determined by multiplying the plasmid copy number with the genome relative abundance. The full set of randomly assigned relative abundances and copy numbers can be found in Supplemental Table 3. Sequences were then concatenated into a single FASTA file with the appropriate relative abundance. MiSeq v3 250bp paired-end reads with a mean fragment length of 1000bp (standard deviation of 50bp) were then simulated using art_illumina (v2016.06.05) [68] resulting in a simulated metagenome of 31,174,411 read pairs. The selection of relative abundance and metagenome simulation itself was performed using the “data_simluation/simulate_metagenome.py” script.

### MAG Recovery

Reads were trimmed using sickle (v1.33) [69] resulting in 25,682,644 surviving read pairs. The trimmed reads were then assembled using 3 different metagenomic assemblers: metaSPAdes (v3.13.0) [18], IDBA-UD (v1.1.3) [19], and megahit (v1.1.3) [20]). The resulting assemblies were summarised using metaQUAST (v5.0.2) [70]. The assemblies were then indexed and reads mapped back using Bowtie 2 (v2.3.4.3) [12].

Samtools (v1.9) was used to sort the read mappings, and the read coverage was calculated using the MetaBAT2 accessory script (jgi_summarize_bam_contig_depths). The three metagenome assemblies were then separately binned using MetaBAT2 (v2.13) [24], and MaxBin 2 (v2.2.6) [25]. MAGs were also recovered using CONCOCT (v0.4.2) [23] following the recommended protocol in the user manual. Briefly, the supplied CONCOCT accessory scripts were used to cut contigs into 10 kilobase fragments (cut_up_fasta.py) and read coverage calculated for the fragments (CONCOCT_coverage_table.py). These fragment coverages were then used to bin the 10kb fragments before the clustered fragments were merged (merge_cutup_clustering.py) to create the final CONCOCT MAG bins (extra_fasta_bins.py). Finally, for each metagenome assembly the predicted bins from these three binners (Maxbin2, MetaBAT 2, and CONCOCT) were combined using the DAS Tool (v1.1.1) meta-binner [26]. This resulted in 12 separate sets of MAGs (one set for each assembler and binner pair).

### MAG assessment

#### Chromosomal Coverage

The MAG assessment for chromosomal coverage was performed by creating a BLASTN 2.9.0+ [71] database consisting of all the chromosomes of the input reference genomes. Each MAG contig was then used as a query against this database and the coverage of the underlying chromosomes tallied by merging the overlapping aligning regions and summing the total length of aligned MAG contigs. The most represented genome in each MAG was assigned as the “identity” of that MAG for further analyses. Coverage values of less than 5% were Altered out and the number of different genomes that contigs from a given MAG aligned to were tallied. Finally, the overall proportion of chromosomes that were not present in any MAG was tallied for each binner and assembler.

In order to investigate the impact of close relatives in the metagenome on ability to bin chromosomes we generated a phylogenetic tree for all the input genomes. Specifically, single copy universal bacterial proteins were identified in the reference genomes using BUSCO v4.0.2 with the Bacteria Odb10 data [72]. The 86 of these proteins that were found in every reference genome were concatenated and aligned using MAFFT v7.427 [73] and masked with trimal v1.4.1-3 [74]. A maximumlikelihood phylogeny was then inferred with IQ-Tree v1.6.12 [75] with the in-built ModelFinder determined partitioning [76]. Pairwise branch distances were then extracted from the resulting tree using ETE3 v3.1.1 [77] and regressed using a linear model against coverage and contamination in seaborn v0.10.0 [78].

#### Plasmid and GI Coverage

Plasmid and GI coverage were assessed in the same way. Firstly, a BLASTN database was generated for each set of MAG contigs. Then each MAG database was searched for plasmid and GI sequences with greater than 50% coverage. All plasmids or GIs which could be found in the unbinned contigs or MAGs were recorded as having been successfully assembled. The subset of these that were found in the binned MAGs was then separately tallied. Finally, we evaluated the proportion of plasmids or GIs that were correctly assigned to the bin that was maximally composed of chromosomes from the same source genome.

### Antimicrobial Resistance and Virulence Factors Assessment

#### Detection of AMR/VF Genes

For the reference genomes, as well as 12 sets of MAGs, prodigal [79] was used to predict open reading frames (ORFs) using the default parameters. AMR genes were predicted using Resistance Gene Identifier (RGI v5.0.0; default parameters) and the Comprehensive Antibiotic Resistance Database (CARD v3.0.3) [80]. Virulence factors were predicted using the predicted ORFs and BLASTX 2.9.0+ [71] against the Virulence Factor Database (VFDB; obtained on Aug 26, 2019) with an e-value cut-off of 0.001 and a minimum identity of 90% [81]. Each MAG was then assigned to a reference chromosome using the above mentioned mapping criteria for downstream analysis.

#### AMR/VF Gene Recovery

For each MAG set, we counted the total number of AMR/VF genes recovered in each metagenomic assembly and each MAG and compared this to the number predicted in their assigned reference chromosome and plasmids. We then assessed the ability for MAGs to correctly bin AMR/VF genes of chromosomal, plasmid, and GI origin by mapping the location of the reference replicon’s predicted genes to the location of the same genes in the MAGs.

#### Protein subcellular localisation predictions

We then sought to assess what the impact of a protein’s predicted subcellular localisation was on its recovery and binning in MAGs. The MAG bins from megahit-DAS Tool assembler-binner combination were selected (as generally best performing) and ORFs predicted using prodigal [79] as above. Subcellular localisation of these proteins were then predicted using PSORTb v3.0 with default parameters and the appropriate Gram setting for that bin’s assigned taxa [82].

## Data availability

All datasets used or generated in this study are available at osf.io/nrejs

## Code availability

All analysis and plotting code used is available at github.com/fmaguire/MAG_gi_plasmid_analysis

## Acknowledgements

This work was supported primarily by a Donald Hill Family Fellowship held by F.M. W.Y.V.L. and B.J. hold Canadian Institutes of Health Research (CIHR) doctoral scholarships. K.G. was supported by a Natural Sciences and Engineering Research Council of Canada (NSERC) Collaborative Research and Training Experience (CREATE) Bioinformatics scholarship. B.J, W.Y.V.L., and K.G. also held Simon Fraser University (SFU) Omics and Data Sciences fellowships. F.S.L.B. holds an SFU Distinguished Professorship and R.G.B. is a Professor and Associate Dean Research at Dalhousie University. Additionally, this work was partially supported by Genome Canada and NSERC grants to R.G.B. and F.S.L.B., and the authors thank the SFU Research Computing Group and Compute Canada for compute resource support.

## Author Information

These authors contributed equally: Finlay Maguire & Baofeng Jia

### Affiliations

*Faculty of Computer Science, Dalhousie University, Halifax, NS, Canada*

Finlay Maguire, & Robert G. Beiko

*Department of Molecular Biology and Biochemistry, Simon Fraser University, Burnaby, BC, Canada*

Baofeng Jia, Kristen Gray, Wing Yin Venus Lau, & Fiona S.L. Brinkman

### Contributions

F.M. and B.J. conceived and designed the study, performed MAG analyses, generated the figures, and wrote the manuscript. W.Y.V.L. and K.G. provided key input on genomic island and subcellular localisation analyses, respectively. F.S.L.B and R.G.B. provided advice throughout the project. All authors contributed to and approved the manuscript.

## Ethics declarations

The authors declare no competing interests.

## Supplementary Information

### Impact of Related Genomes on MAG

By generating a phylogeny of universal single copy genes in our input genomes we analysed the relationship between the presence of closely related genomes and the ability of the different MAG-recovery methods to bin chromosomal sequences. Specifically, we regressed phylogenetic distance on this phylogeny with per-bin chromosomal coverage (Fig. S1) and bin purity (Fig. S2). This identified no clear relationship between chromosomal coverage and the phylogenetic distance to the nearest relative in the metagenome (Fig. S1), however, there did seem to be a negative correlation between phylogenetic distance to closest relative and the purity of a MAG bin (Fig. S2). In other words, across all methods, a MAG bin was more likely to have multiple genomes present if there were close relatives.

**Figure S1:**
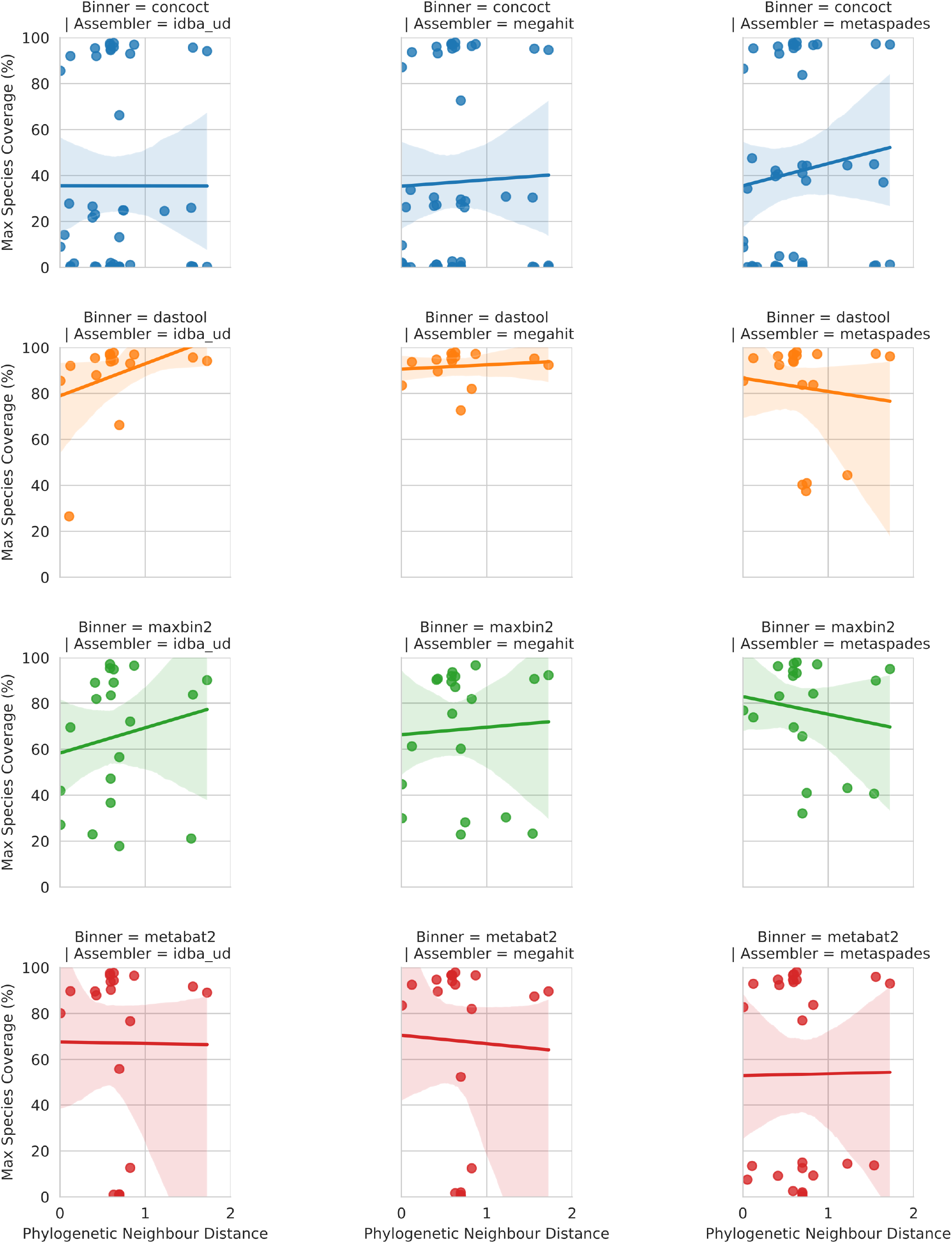
Relationship between phylogenetic distance to closest neighbour input genome on genomic coverage in MAG majority comprised of that taxa. Each dot represents the genomic coverage of a particular taxa and the branch distance on an 86-protein concatenated phylogeny between that taxa and its nearest neighbour. Rows indicate the binning software and columns the metagenomic assembler. Regression line is a simple linear model fitted in seaborn.

**Figure S2:**
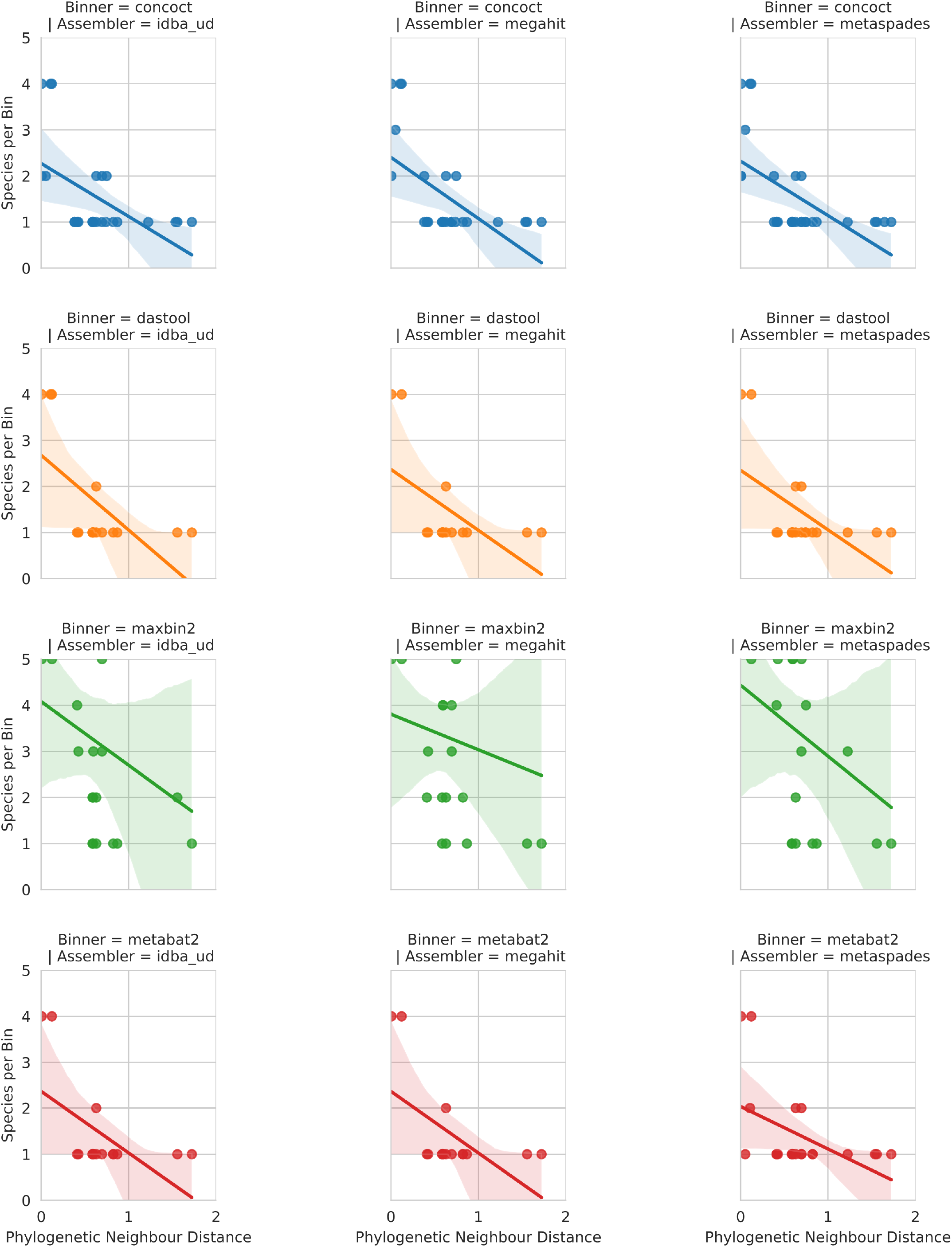
Relationship between phylogenetic distance to closest neighbour input genome on bin purity. Each dot shows the number of other input genomes detectable in a given MAG bin in relation to the branch distance on an 86-protein concatenated phylogeny between the majority taxa in that bin and its nearest neighbour.

### Recovery of Specific Gene Content

We explored the ability of different approaches to find open reading frames (ORFs) within MAGs. Overall, the total number of predicted ORFs in MAGs followed a similar trend (Fig. S3) as the chromosomal coverage and purity (Fig. 2). Of the four binning tools, CONCOCT performed the worst, finding <30% of the number of ORFs in our reference genomes used to construct the synthetic data. MetaBAT2 performed second worst at ~80%. DASTool recovered a similar number to our reference and Maxbin2 detected 7-46% more genes. The Assembler method did not significantly impact the number of genes predicted with the exception of Maxbin2, in which IDBA_UD was the closest to reference and metaSPAdes predicted 46% more ORFs. Given that there is reason to suspect that there are some issues with the ORF calling in the MAGs. i.e. some tools produced more predicted ORFs than reference, it could be the case that some of these sequences are present in the assemblies (with errors/gaps), but are not being identified as ORFs, or are broken into multiple ORFs, leading to issues downstream labeling them correctly as AMR/VF genes. Regardless of different tools producing a different number of ORFs, the recovery of AMR/VF is pretty consistent regardless of how many ORFs are predicted.

**Figure S3:**
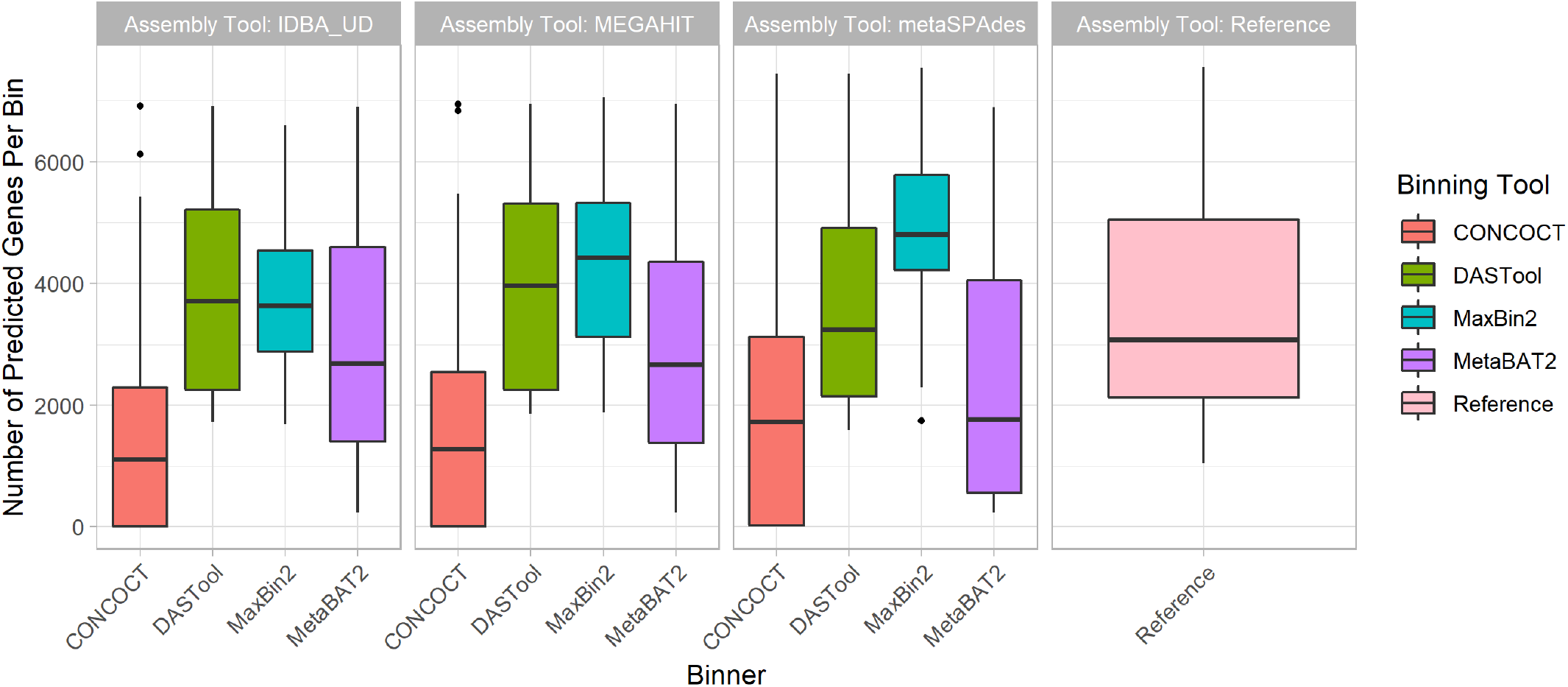
Predicted Gene Content. The total number of open reading frames (ORF) predicted followed the same trend as chromosomal coverage and purity. The assemblers (colored bars) did not contribute to variability in the number of ORFs detected. Of the 4 binners, CONCOCT recovered <30% of our reference genome ORFs. DASTool and MetaBAT2 predicted a similar number as our reference genomes.

### Comparisons of Rates of Loss

Combining the performance metrics for Figs. 3, 4, 5, and 6 to compare the rates of loss of different components emphasises some of the observed patterns (see Fig. S4). This highlights that genomic components (GIs and plasmids) and plasmids in particular are lost at a higher rate than individual gene types during MAG recovery.

**Figure S4:**
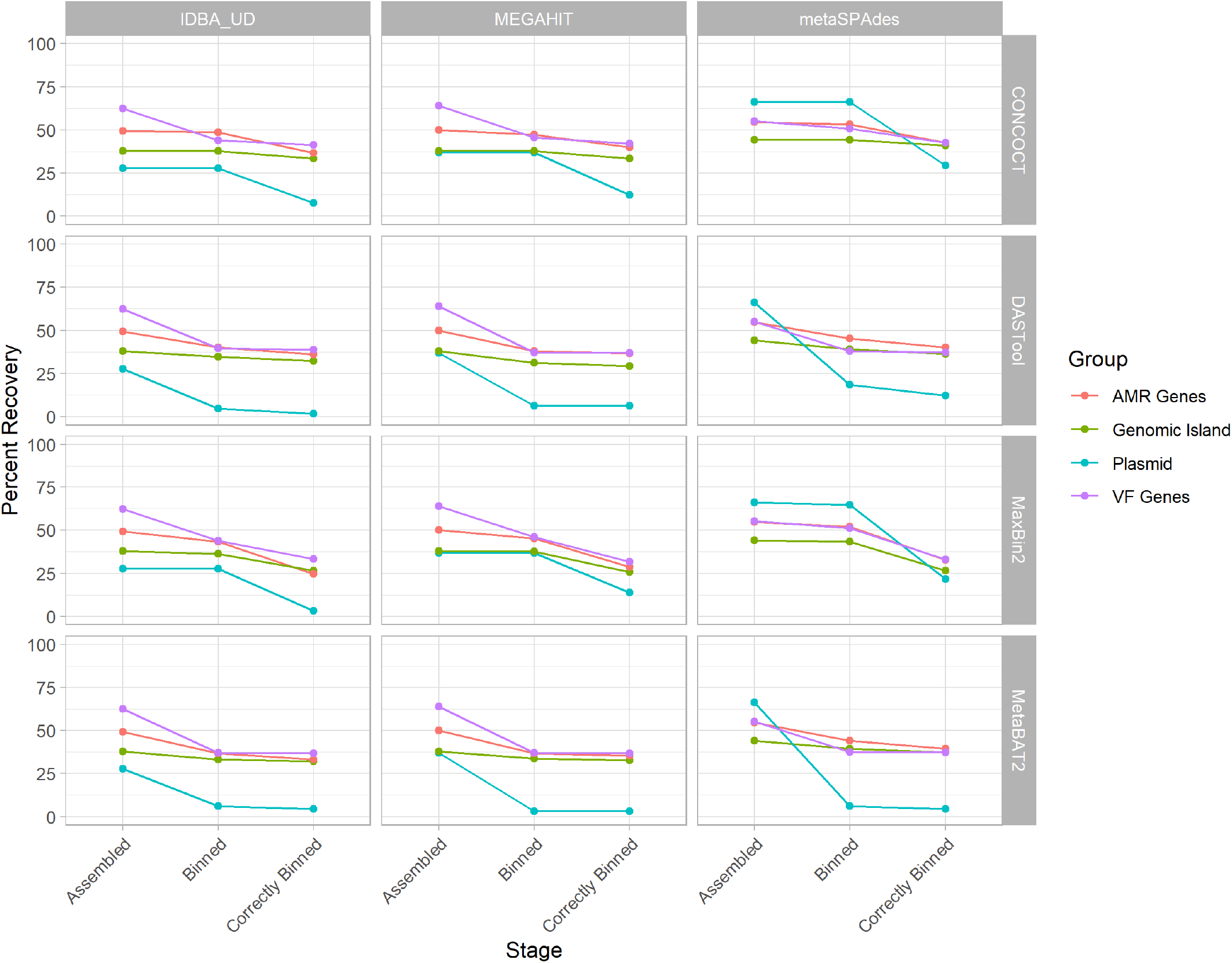
Comparison of rates of loss for different genomic components and gene types across assemblers and binning tools. Each line represents a different component as indicated by the legend with assemblers indicated by row and binning tool by column. This shows that regardless of approach genomic components (GIs and plasmids) are lost at a higher rate than individual VF or AMR genes.

## References

1. Genomic analysis of uncultured marine viral communities M. Breitbart, P. Salamon, B. Andresen, J. M. Mahaffy, A. M. Segall, D. Mead, F. Azam, F. Rohwer Proceedings of the National Academy of Sciences (2002-10-16) https://doi.org/br7jq3 DOI: 10.1073/pnas.202488399 · PMID: 12384570 · PMCID: PMC137870

2. Shotgun metagenomics, from sampling to analysis Christopher Quince, Alan W Walker, Jared T Simpson, Nicholas J Loman, Nicola Segata Nature Biotechnology (2017-09-12) https://doi.org/gbv6nf DOI: 10.1038/nbt.3935 · PMID: 28898207

3. A Systematic Analysis of Biosynthetic Gene Clusters in the Human Microbiome Reveals a Common Family of Antibiotics Mohamed S. Donia, Peter Cimermancic, Christopher J. Schulze, Laura C. Wieland Brown, John Martin, Makedonka Mitreva, Jon Clardy, Roger G. Linington, Michael A. Fischbach Cell (2014-09) https://doi.org/f6k3fg DOI: 10.1016/j.cell.2014.08.032 · PMID: 25215495 · PMCID: PMC4164201

4. Expanding the soil antibiotic resistome: exploring environmental diversity Vanessa M D’Costa, Emma Griffiths, Gerard D Wright Current Opinion in Microbiology (2007-10) https://doi.org/cfbpjj DOI: 10.1016/j.mib.2007.08.009 · PMID: 17951101

5. Antibiotic resistance is ancient Vanessa M. D’Costa, Christine E. King, Lindsay Kalan, Mariya Morar, Wilson W. L. Sung, Carsten Schwarz, Duane Froese, Grant Zazula, Fabrice Calmels, Regis Debruyne,… Gerard D. Wright Nature (2011-08-31) https://doi.org/b3wbvx DOI: 10.1038/nature10388 · PMID: 21881561

6. A Culture-Independent Sequence-Based Metagenomics Approach to the Investigation of an Outbreak of Shiga-Toxigenic Escherichia coli O104:H4 Nicholas J. Loman, Chrystala Constantinidou, Martin Christner, Holger Rohde, Jacqueline Z.-M. Chan, Joshua Quick, Jacqueline C. Weir, Christopher Quince, Geoffrey P. Smith, Jason R. Betley,… Mark J. Pallen JAMA (2013-04-10) https://doi.org/f5rqft DOI: 10.1001/jama.2013.3231 · PMID: 23571589

7. A first look at the Oxford Nanopore MinION sequencer Alexander S. Mikheyev, Mandy M. Y. Tin Molecular Ecology Resources (2014-11) https://doi.org/vmt DOI: 10.1111/1755-0998.12324 · PMID: 25187008

8. Real-Time DNA Sequencing from Single Polymerase Molecules J. Eid, A. Fehr, J. Gray, K. Luong, J. Lyle, G. Otto, P. Peluso, D. Rank, P. Baybayan, B. Bettman,… S. Turner Science (2009-01-02) https://doi.org/cz7ndk DOI: 10.1126/science.1162986 · PMID: 19023044

9. Ultra-deep, long-read nanopore sequencing of mock microbial community standards Samuel M Nicholls,Joshua C Quick, Shuiquan Tang, Nicholas J Loman GigaScience (2019-05) https://doi.org/gf39g3 DOI: 10.1093/gigascience/giz043 · PMID: 31089679 · PMCID: PMC6520541

10. Long-read based de novo assembly of low-complexity metagenome samples results in finished genomes and reveals insights into strain diversity and an active phage system Vincent Somerville, Stefanie Lutz, Michael Schmid, Daniel Frei, Aline Moser, Stefan Irmler, Jürg E. Frey, Christian H. Ahrens BMC Microbiology (2019-06-25) https://doi.org/gf5ffc DOI: 10.1186/s12866-019-1500-0 · PMID: 31238873 · PMCID: PMC6593500

11. Fast and sensitive protein alignment using DIAMOND Benjamin Buchfink, Chao Xie, Daniel H Huson Nature Methods (2014-11-17) https://doi.org/gftzcs DOI: 10.1038/nmeth.3176 · PMID: 25402007

12. Fast gapped-read alignment with Bowtie 2 Ben Langmead, Steven L Salzberg Nature Methods (2012-03-04) https://doi.org/gd2xzn DOI: 10.1038/nmeth.1923 · PMID: 22388286 · PMCID: PMC3322381

13. nhmmer: DNA homology search with profile HMMs T. J. Wheeler, S. R. Eddy Bioinformatics (2013-07-09) https://doi.org/f5xm9x DOI: 10.1093/bioinformatics/btt403 · PMID: 23842809 · PMCID: PMC3777106

14. CLARK: fast and accurate classification of metagenomic and genomic sequences using discriminative k-mers Rachid Ounit, Steve Wanamaker, Timothy J Close, Stefano Lonardi BMC Genomics (2015-03-25) https://doi.org/gb3h2t DOI: 10.1186/s12864-015-1419-2 · PMID: 25879410 · PMCID: PMC4428112

15. vanM, a New Glycopeptide Resistance Gene Cluster Found in Enterococcus faecium X. Xu, D. Lin, G. Yan, X. Ye, S. Wu, Y. Guo, D. Zhu, F. Hu, Y. Zhang, F. Wang,… M. Wang Antimicrobial Agents and Chemotherapy (2010-08-23) https://doi.org/cnpst5 DOI: 10.1128/aac.01710-09 · PMID: 20733041 · PMCID: PMC2976141

16. Co-selection of antibiotic and metal resistance Craig Baker-Austin, Meredith S. Wright, Ramunas Stepanauskas, J. V. McArthur Trends in Microbiology (2006-04) https://doi.org/fvkg6d DOI: 10.1016/j.tim.2006.02.006 · PMID: 16537105

17. Gene flow, mobile genetic elements and the recruitment of antibiotic resistance genes into Gram-negative pathogens Hatch W. Stokes, Michael R. Gillings FEMS Microbiology Reviews (2011-09) https://doi.org/fw543p DOI: 10.1111/j.1574-6976.2011.00273.x · PMID: 21517914

18. metaSPAdes: a new versatile metagenomic assembler Sergey Nurk, Dmitry Meleshko, Anton Korobeynikov, Pavel A. Pevzner Genome Research (2017-05) https://doi.org/f97jkv DOI: 10.1101/gr.213959.116 · PMID: 28298430 · PMCID: PMC5411777

19. IDBA-UD: a de novo assembler for single-cell and metagenomic sequencing data with highly uneven depth Y. Peng, H. C. M. Leung, S. M. Yiu, F. Y. L. Chin Bioinformatics (2012-04-11) https://doi.org/f3z7hv DOI: 10.1093/bioinformatics/bts174 · PMID: 22495754

20. MEGAHIT: an ultra-fast single-node solution for large and complex metagenomics assembly via succinct de Bruijn graph Dinghua Li, Chi-Man Liu, Ruibang Luo, Kunihiko Sadakane, Tak-Wah Lam Bioinformatics (2015-05-15) https://doi.org/f7fb5z DOI: 10.1093/bioinformatics/btv033 · PMID: 25609793

21. Community structure and metabolism through reconstruction of microbial genomes from the environment Gene W. Tyson, Jarrod Chapman, Philip Hugenholtz, Eric E. Allen, Rachna J. Ram, Paul M. Richardson, Victor V. Solovyev, Edward M. Rubin, Daniel S. Rokhsar, Jillian F. Banfield Nature (2004-02-01) https://doi.org/b85j5j DOI: 10.1038/nature02340 · PMID: 14961025

22. A review of methods and databases for metagenomic classification and assembly Florian P Breitwieser,Jennifer Lu, Steven L Salzberg Briefings in Bioinformatics (2019-07) https://doi.org/gdq95k DOI: 10.1093/bib/bbx120 · PMID: 29028872 · PMCID: PMC6781581

23. COCACOLA: binning metagenomic contigs using sequence COmposition, read CoverAge, COalignment and paired-end read LinkAge Yang Young Lu,Ting Chen, Jed A. Fuhrman, Fengzhu Sun Bioinformatics (2016-06-02) https://doi.org/f9x7sc DOI: 10.1093/bioinformatics/btw290 · PMID: 27256312

24. MetaBAT 2: an adaptive binning algorithm for robust and efficient genome reconstruction from metagenome assemblies Dongwan Kang, Feng Li, Edward S Kirton, Ashleigh Thomas, Rob S Egan, Hong An, Zhong Wang (2019-02-06) https://doi.org/gf5fhv DOI: 10.7287/peerj.preprints.27522v1

25. MaxBin 2.0: an automated binning algorithm to recover genomes from multiple metagenomic datasets Yu-Wei Wu, Blake A. Simmons, Steven W. Singer Bioinformatics (2016-02-15) https://doi.org/f8c9n2 DOI: 10.1093/bioinformatics/btv638 · PMID: 26515820

26. Recovery of genomes from metagenomes via a dereplication, aggregation and scoring strategy Christian M. K. Sieber, Alexander J. Probst, Allison Sharrar, Brian C. Thomas, Matthias Hess, Susannah G. Tringe, Jillian F. Banfield Nature Microbiology (2018-05-28) https://doi.org/gfwwfg DOI: 10.1038/s41564-018-0171-1 · PMID: 29807988 · PMCID: PMC6786971

27. Unusual biology across a group comprising more than 15% of domain Bacteria Christopher T. Brown, Laura A. Hug, Brian C. Thomas, Itai Sharon, Cindy J. Castelle, Andrea Singh, Michael J. Wilkins, Kelly C. Wrighton, Kenneth H. Williams, Jillian F. Banfield Nature (2015-06-15) https://doi.org/f7h5xj DOI: 10.1038/nature14486 · PMID: 26083755

28. Recovery of nearly 8,000 metagenome-assembled genomes substantially expands the tree of life Donovan H. Parks, Christian Rinke, Maria Chuvochina, Pierre-Alain Chaumeil, Ben J. Woodcroft, Paul N. Evans, Philip Hugenholtz, Gene W. Tyson Nature Microbiology (2017-09-11) https://doi.org/cczd DOI: 10.1038/s41564-017-0012-7 · PMID: 28894102

29. The genomic and proteomic landscape of the rumen microbiome revealed by comprehensive genome-resolved metagenomics Robert D. Stewart, Marc D. Auffret, Amanda Warr, Alan W. Walker, Rainer Roehe, Mick Watson bioRxiv (2018-12-08) https://doi.org/gf5fhr DOI: 10.1101/489443

30. Genome-centric view of carbon processing in thawing permafrost Ben J. Woodcroft, Caitlin M. Singleton, Joel A. Boyd, Paul N. Evans, Joanne B. Emerson, Ahmed A. F. Zayed, Robert D. Hoelzle, Timothy O. Lamberton, Carmody K. McCalley, Suzanne B. Hodgkins,… Gene W. Tyson Nature (2018-07-16) https://doi.org/gdth6p DOI: 10.1038/s41586-018-0338-1 · PMID: 30013118

31. Mediterranean grassland soil C–N compound turnover is dependent on rainfall and depth, and is mediated by genomically divergent microorganisms Spencer Diamond, Peter F. Andeer, Zhou Li, Alexander Crits-Christoph, David Burstein, Karthik Anantharaman, Katherine R. Lane, Brian C. Thomas, Chongle Pan, Trent R. Northen, Jillian F. Banfield Nature Microbiology (2019-05-20) https://doi.org/gf5fcx DOI: 10.1038/s41564-019-0449-y · PMID: 31110364 · PMCID: PMC6784897

32. AMBER: Assessment of Metagenome BinnERs Fernando Meyer, Peter Hofmann, Peter Belmann, Ruben Garrido-Oter, Adrian Fritz, Alexander Sczyrba, Alice C McHardy GigaScience (2018-06) https://doi.org/gdptz9 DOI: 10.1093/gigascience/giy069 · PMID: 29893851 · PMCID: PMC6022608

33. Detecting genomic islands using bioinformatics approaches Morgan G. I. Langille, William W. L. Hsiao, Fiona S. L. Brinkman Nature Reviews Microbiology (2010-05) https://doi.org/d6ss55 DOI: 10.1038/nrmicro2350 · PMID: 20395967

34. Horizontal gene transfer: building the web of life Shannon M. Soucy, Jinling Huang, Johann Peter Gogarten Nature Reviews Genetics(2015-07-17) https://doi.org/f7j3d9 DOI: 10.1038/nrg3962 · PMID: 26184597

35. The Association of Virulence Factors with Genomic Islands Shannan J. Ho Sui, Amber Fedynak, William W. L. Hsiao, Morgan G. I. Langille, Fiona S. L. Brinkman PLoS ONE (2009-12-01) https://doi.org/c7hsvv DOI: 10.1371/journal.pone.0008094 · PMID: 19956607 · PMCID: PMC2779486

36. Dissemination of Antimicrobial Resistance in Microbial Ecosystems through Horizontal Gene Transfer Christian J. H. von Wintersdorff, John Penders, Julius M. van Niekerk, Nathan D. Mills, Snehali Majumder, Lieke B. van Alphen, Paul H. M. Savelkoul, Petra F. G. Wolffs Frontiers in Microbiology (2016-02-19) https://doi.org/gf5fht DOI: 10.3389/fmicb.2016.00173 · PMID: 26925045 · PMCID: PMC4759269

37. T ransfer of antibiotic-resistance genes via phage-related mobile elements Maryury Brown-Jaque, William Calero-Cáceres, Maite Muniesa Plasmid (2015-05) https://doi.org/f7dvxv DOI: 10.1016/j.plasmid.2015.01.001 · PMID: 25597519

38. SIGI: score-based identification of genomic islands. Rainer Merkl BMC bioinformatics (2004-03-03) https://www.ncbi.nlm.nih.gov/pubmed/15113412 DOI: 10.1186/1471-2105-5-22 · PMID: 15113412 · PMCID: PMC394314

39. Improved genomic island predictions with IslandPath-DIMOB Claire Bertelli, Fiona SL Brinkman Bioinformatics (2018-07-01) https://doi.org/gdphgs DOI: 10.1093/bioinformatics/bty095 · PMID: 29905770 · PMCID: PMC6022643

40. IslandViewer 3: more flexible, interactive genomic island discovery, visualization and analysis: Figure 1. Bhavjinder K. Dhillon, Matthew R. Laird, Julie A. Shay, Geoffrey L. Winsor, Raymond Lo, Fazmin Nizam, Sheldon K. Pereira, Nicholas Waglechner, Andrew G. McArthur, Morgan G. I. Langille, Fiona S. L. Brinkman Nucleic Acids Research (2015-07-01) https://doi.org/f7n2xs DOI: 10.1093/nar/gkv401 · PMID: 25916842 · PMCID: PMC4489224

41. Microbial genomic island discovery, visualization and analysis Claire Bertelli, Keith E Tilley, Fiona SL Brinkman Briefings in Bioinformatics (2019-09) https://doi.org/gdnhfv DOI: 10.1093/bib/bby042 · PMID: 29868902 · PMCID: PMC6917214

42. Multicopy plasmids potentiate the evolution of antibiotic resistance in bacteria Alvaro San Millan,Jose Antonio Escudero, Danna R. Gifford, Didier Mazel, R. Craig MacLean Nature Ecology & Evolution (2016-11-07) https://doi.org/bs76 DOI: 10.1038/s41559-016-0010 · PMID: 28812563

43. Small-Plasmid-Mediated Antibiotic Resistance Is Enhanced by Increases in Plasmid Copy Number and Bacterial Fitness Alvaro San Millan,Alfonso Santos-Lopez, Rafael Ortega-Huedo, Cristina Bernabe-Balas, Sean P. Kennedy, Bruno Gonzalez-Zorn Antimicrobial Agents and Chemotherapy (2015-06) https://doi.org/f7k8bk DOI: 10.1128/aac.00235-15 · PMID: 25824216 · PMCID: PMC4432117

44. cBar: a computer program to distinguish plasmid-derived from chromosome-derived sequence fragments in metagenomics data Fengfeng Zhou, Ying Xu Bioinformatics (2010-08-15) https://doi.org/cn7486 DOI: 10.1093/bioinformatics/btq299 · PMID: 20538725 · PMCID: PMC2916713

45. Modal Codon Usage: Assessing the Typical Codon Usage of a Genome J. J. Davis, G. J. Olsen Molecular Biology and Evolution (2009-12-17) https://doi.org/bhsmq5 DOI: 10.1093/molbev/msp281 · PMID: 20018979 · PMCID: PMC2839124

46. Understanding the mechanisms and drivers of antimicrobial resistance. Alison H Holmes, Luke SP Moore, Arnfinn Sundsfjord, Martin Steinbakk, Sadie Regmi, Abhilasha Karkey, Philippe J Guerin, Laura JV Piddock Lancet (London, England) (2015-11-18) https://www.ncbi.nlm.nih.gov/pubmed/26603922 DOI: 10.1016/s0140-6736(15)00473-0 · PMID: 26603922

47. On the (im)possibility of reconstructing plasmids from whole-genome short-read sequencing data Sergio Arredondo-Alonso, Rob J. Willems, Willem van Schaik, Anita C. Schürch Microbial Genomics (2017-10-01) https://doi.org/gf6b63 DOI: 10.1099/mgen.0.000128 · PMID: 29177087 · PMCID: PMC5695206

48. SCAPP: An algorithm for improved plasmid assembly in metagenomes David Pellow, Maraike Probst, Ori Furman, Alvah Zorea, Arik Segal, Itzik Mizrahi, Ron Shamir bioRxiv (2020-01-14) https://doi.org/ggkt4f DOI: 10.1101/2020.01.12.903252

49. Complete and validated genomes from a metagenome Daniel J Giguere,Alexander T Bahcheli, Benjamin R Joris, Julie M Paulssen, Lisa M Gieg, Martin W Flatley, Gregory B Gloor bioRxiv (2020-04-09) https://doi.org/ggsnqn DOI: 10.1101/2020.04.08.032540

50. Long-read metagenomic exploration of extrachromosomal mobile genetic elements in the human gut Yoshihiko Suzuki, Suguru Nishijima, Yoshikazu Furuta, Jun Yoshimura, Wataru Suda, Kenshiro Oshima, Masahira Hattori, Shinichi Morishita Microbiome (2019-08-27) https://doi.org/ggsnqp DOI: 10.1186/s40168-019-0737-z · PMID: 31455406 · PMCID: PMC6712665

51. Loss of microbial diversity and pathogen domination of the gut microbiota in critically ill patients Anuradha Ravi, Fenella D Halstead, Amy Bamford, Anna Casey, Nicholas M. Thomson, Willem van Schaik, Catherine Snelson, Robert Goulden, Ebenezer Foster-Nyarko, George M. Savva,… Beryl A. Oppenheim Microbial Genomics (2019-09-01) https://doi.org/dcc9 DOI: 10.1099/mgen.0.000293 · PMID: 31526447 · PMCID: PMC6807385

52. Metagenomic and metatranscriptomic analyses reveal activity and hosts of antibiotic resistance genes in activated sludge Zongbao Liu, Uli Klümper, Yang Liu, Yuchun Yang, Qiaoyan Wei, Jih-Gaw Lin, Ji-Dong Gu, Meng Li Environment International (2019-08) https://doi.org/ggsqc9 DOI: 10.1016/j.envint.2019.05.036 · PMID: 31129497

53. Genome-resolved metagenomics to study co-occurrence patterns and intraspecific heterogeneity among plant pathogen metapopulations Eric Newberry, Rishi Bhandari, Joseph Kemble, Edward Sikora, Neha Potnis Environmental Microbiology (2020-04-02) https://doi.org/ggsqdd DOI: 10.1111/1462-2920.14989 · PMID: 32207218

54. Benefits of Genomic Insights and CRISPR-Cas Signatures to Monitor Potential Pathogens across Drinking Water Production and Distribution Systems Ya Zhang, Masaaki Kitajima, Andrew J. Whittle, Wen-Tso Liu Frontiers in Microbiology (2017-10-19) https://doi.org/gcpmqk DOI: 10.3389/fmicb.2017.02036 · PMID: 29097994 · PMCID: PMC5654357

55. Metagenomics of Two Severe Foodborne Outbreaks Provides Diagnostic Signatures and Signs of Coinfection Not Attainable by Traditional Methods Andrew D. Huang, Chengwei Luo, Angela Pena-Gonzalez, Michael R. Weigand, Cheryl L. Tarr, Konstantinos T. Konstantinidis Applied and Environmental Microbiology (2016-11-23) https://doi.org/ggsqdf DOI: 10.1128/aem.02577-16 · PMID: 27881416 · PMCID: PMC5244306

56. Gene fragmentation in bacterial draft genomes: extent, consequences and mitigation Jonathan L Klassen,Cameron R Currie BMC Genomics (2012) https://doi.org/fzg6gg DOI: 10.1186/1471-2164-13-14 · PMID: 22233127 · PMCID: PMC3322347

57. Detection of bacterial pathogens from clinical specimens using conventional microbial culture and 16S metagenomics: a comparative study Lalanika M. Abayasekara, Jennifer Perera, Vishvanath Chandrasekharan, Vaz S. Gnanam, Nisala A. Udunuwara, Dileepa S. Liyanage, Nuwani E. Bulathsinhala, Subhashanie Adikary, Janith V. S. Aluthmuhandiram, Chrishanthi S. Thanaseelan,… Janahan Ilango BMC Infectious Diseases (2017-09-19) https://doi.org/ggsr88 DOI: 10.1186/s12879-017-2727-8 · PMID: 28927397 · PMCID: PMC5606128

58. Characterization of Bacterial Community Diversity in Cystic Fibrosis Lung Infections by Use of 16S Ribosomal DNA Terminal Restriction Fragment Length Polymorphism Profiling G. B. Rogers, M. P. Carroll, D. J. Serisier, P. M. Hockey, G. Jones, K. D. Bruce Journal of Clinical Microbiology (2004-11-04) https://doi.org/ctw2bk DOI: 10.1128/jcm.42.11.5176-5183.2004 · PMID: 15528712 · PMCID: PMC525137

59. The vaginal microbiome of pregnant women is less rich and diverse, with lower prevalence of Mollicutes, compared to non-pregnant women Aline C. Freitas, Bonnie Chaban, Alan Bocking, Maria Rocco, Siwen Yang, Janet E. Hill, Deborah M. Money, The VOGUE Research Group Scientific Reports (2017-08-23) https://doi.org/gbtz3z DOI: 10.1038/s41598-017-07790-9 · PMID: 28835692 · PMCID: PMC5569030

60. 16S rDNA Pyrosequencing Analysis of Bacterial Community in Heavy Metals Polluted Soils Marcin Gołębiewski, Edyta Deja-Sikora, Marcin Cichosz, Andrzej Tretyn, Borys Wróbel Microbial Ecology (2014-01-09) https://doi.org/ggsr86 DOI: 10.1007/s00248-013-0344-7 · PMID: 24402360 · PMCID: PMC3962847

61. Comparison of Species Richness Estimates Obtained Using Nearly Complete Fragments and Simulated Pyrosequencing-Generated Fragments in 16S rRNA Gene-Based Environmental Surveys N. Youssef, C. S. Sheik, L. R. Krumholz, F. Z. Najar, B. A. Roe, M. S. Elshahed Applied and Environmental Microbiology (2009-06-26) https://doi.org/chd3zf DOI: 10.1128/aem.00592-09 · PMID: 19561178 · PMCID: PMC2725448

62. Comparative Analysis of Pyrosequencing and a Phylogenetic Microarray for Exploring Microbial Community Structures in the Human Distal Intestine Marcus J. Claesson, Orla O’Sullivan, Qiong Wang, Janne Nikkilä, Julian R. Marchesi, Hauke Smidt, Willem M. de Vos, R. Paul Ross, Paul W. O’Toole PLoS ONE (2009-08-20) https://doi.org/fqn5td DOI: 10.1371/journal.pone.0006669 · PMID: 19693277 · PMCID: PMC2725325

63. Metagenomic characterization of the effect of feed additives on the gut microbiome and antibiotic resistome of feedlot cattle Milton Thomas, Megan Webb, Sudeep Ghimire, Amanda Blair, Kenneth Olson, Gavin John Fenske, Alex Thomas Fonder, Jane Christopher-Hennings, Derek Brake, Joy Scaria Scientific Reports (2017-09-25) https://doi.org/gb26tf DOI: 10.1038/s41598-017-12481-6 · PMID: 28947833 · PMCID: PMC5612972

64. CAMISIM: simulating metagenomes and microbial communities Adrian Fritz, Peter Hofmann, Stephan Majda, Eik Dahms, Johannes Dröge, Jessika Fiedler, Till R. Lesker, Peter Belmann, Matthew Z. DeMaere, Aaron E. Darling,… Alice C. McHardy Microbiome (2019-02-08) https://doi.org/ggrncx DOI: 10.1186/s40168-019-0633-6 · PMID: 30736849 · PMCID: PMC6368784

65. mockrobiota: a Public Resource for Microbiome Bioinformatics Benchmarking Nicholas A. Bokulich, Jai Ram Rideout, William G. Mercurio, Arron Shiffer, Benjamin Wolfe, Corinne F. Maurice, Rachel J. Dutton, Peter J. Turnbaugh, Rob Knight, J. Gregory Caporaso mSystems (2016-10-18) https://doi.org/gbrmqs DOI: 10.1128/msystems.00062-16 · PMID: 27822553 · PMCID: PMC5080401

66. ARIBA: rapid antimicrobial resistance genotyping directly from sequencing reads Martin Hunt, Alison E Mather, Leonor Sánchez-Busó, Andrew J Page, Julian Parkhill, Jacqueline A Keane, Simon R Harris Microbial Genomics (2017-10-01) https://doi.org/gf5fd9 DOI: 10.1099/mgen.0.000131 · PMID: 29177089 · PMCID: PMC5695208

67. Critical Assessment of Metagenome Interpretation—a benchmark of metagenomics software Alexander Sczyrba, Peter Hofmann, Peter Belmann, David Koslicki, Stefan Janssen, Johannes Dröge, Ivan Gregor, Stephan Majda, Jessika Fiedler, Eik Dahms,… Alice C McHardy Nature Methods (2017-10-02) https://doi.org/gbzspt DOI: 10.1038/nmeth.4458 · PMID: 28967888 · PMCID: PMC5903868

68. ART: a next-generation sequencing read simulator Weichun Huang, Leping Li, Jason R. Myers, Gabor T. Marth Bioinformatics (2012-02-15) https://doi.org/fzf84c DOI: 10.1093/bioinformatics/btr708 · PMID: 22199392 · PMCID: PMC3278762

69. Sickle: A sliding-window, adaptive, quality-based trimming tool for FastQ files NA Joshi, JN Fass GitHub (2011) https://github.com/najoshi/sickle

70. MetaQUAST: evaluation of metagenome assemblies Alla Mikheenko, Vladislav Saveliev, Alexey Gurevich Bioinformatics (2016-04-01) https://doi.org/f8jdjj DOI: 10.1093/bioinformatics/btv697 · PMID: 26614127

71. BLAST+: architecture and applications Christiam Camacho, George Coulouris, Vahram Avagyan, Ning Ma, Jason Papadopoulos, Kevin Bealer, Thomas L Madden BMC Bioinformatics (2009) https://doi.org/cnjxgz DOI: 10.1186/1471-2105-10-421 · PMID: 20003500 · PMCID: PMC2803857

72. BUSCO: assessing genome assembly and annotation completeness with single-copy orthologs Felipe A. Simão, Robert M. Waterhouse, Panagiotis Ioannidis, Evgenia V. Kriventseva, Evgeny M. Zdobnov Bioinformatics (2015-10-01) https://doi.org/gfznpw DOI: 10.1093/bioinformatics/btv351 · PMID: 26059717

73. Parallelization of MAFFT for large-scale multiple sequence alignments Tsukasa Nakamura, Kazunori D Yamada, Kentaro Tomii, Kazutaka Katoh Bioinformatics (2018-07-15) https://doi.org/gc4th3 DOI: 10.1093/bioinformatics/bty121 · PMID: 29506019 · PMCID: PMC6041967

74. trimAl: a tool for automated alignment trimming in large-scale phylogenetic analyses S. Capella-Gutierrez, J. M. Silla-Martinez, T. Gabaldon Bioinformatics (2009-06-08) https://doi.org/bjhdh7 DOI: 10.1093/bioinformatics/btp348 · PMID: 19505945 · PMCID: PMC2712344

75. IQ-TREE: A Fast and Effective Stochastic Algorithm for Estimating Maximum-Likelihood Phylogenies Lam-Tung Nguyen, Heiko A. Schmidt, Arndt von Haeseler, Bui Quang Minh Molecular Biology and Evolution (2015-01) https://doi.org/f3srtd DOI: 10.1093/molbev/msu300 · PMID: 25371430 · PMCID: PMC4271533

76. PartitionFinder: Combined Selection of Partitioning Schemes and Substitution Models for Phylogenetic Analyses R. Lanfear, B. Calcott, S. Y. W. Ho, S. Guindon Molecular Biology and Evolution (2012-01-20) https://doi.org/fzgsw3 DOI: 10.1093/molbev/mss020 · PMID: 22319168

77. ETE 3: Reconstruction, Analysis, and Visualization of Phylogenomic Data Jaime Huerta-Cepas, François Serra, Peer Bork Molecular Biology and Evolution (2016-06) https://doi.org/gfzpph DOI: 10.1093/molbev/msw046 · PMID: 26921390 · PMCID: PMC4868116

78. mwaskom/seaborn: v0.10.0 (January 2020) Michael Waskom, Olga Botvinnik, Joel Ostblom, Saulius Lukauskas, Paul Hobson, MaozGelbart, David C Gemperline, Tom Augspurger, Yaroslav Halchenko, John B. Cole,… Constantine Evans Zenodo (2020-01-24) https://doi.org/ggkff7 DOI: 10.5281/zenodo.3629446

79. Prodigal: prokaryotic gene recognition and translation initiation site identification Doug Hyatt, Gwo-Liang Chen, Philip F LoCascio, Miriam L Land, Frank W Larimer, Loren J Hauser BMC Bioinformatics (2010-03-08) https://doi.org/cktxnm DOI: 10.1186/1471-2105-11-119 · PMID: 20211023 · PMCID: PMC2848648

80. CARD 2020: antibiotic resistome surveillance with the comprehensive antibiotic resistance database Brian P Alcock,Amogelang R Raphenya, Tammy TY Lau, Kara K Tsang, Mégane Bouchard, Arman Edalatmand, William Huynh, Anna-Lisa V Nguyen, Annie A Cheng, Sihan Liu,… Andrew G McArthur Nucleic Acids Research (2019-10-29) https://doi.org/ggckg6 DOI: 10.1093/nar/gkz935 · PMID: 31665441 · PMCID: PMC7145624

81. VFDB 2019: a comparative pathogenomic platform with an interactive web interface Bo Liu, Dandan Zheng, Qi Jin, Lihong Chen, Jian Yang Nucleic Acids Research (2019-01-08) https://doi.org/gf4zfr DOI: 10.1093/nar/gky1080 · PMID: 30395255 · PMCID: PMC6324032

82. PSORTb 3.0: improved protein subcellular localization prediction with refined localization subcategories and predictive capabilities for all prokaryotes Nancy Y. Yu, James R. Wagner, Matthew R. Laird, Gabor Melli, Sébastien Rey, Raymond Lo, Phuong Dao, S. Cenk Sahinalp, Martin Ester, Leonard J. Foster, Fiona S. L. Brinkman Bioinformatics (2010-07-01) https://doi.org/bz3q2w DOI: 10.1093/bioinformatics/btq249 · PMID: 20472543 · PMCID: PMC2887053

